# Essential role for HSP40 in asexual replication and thermotolerance of malaria parasites

**DOI:** 10.1101/2024.11.05.622024

**Authors:** Brianne Roper, Deepika Kannan, Emily S. Mathews, Audrey R. Odom John

## Abstract

*Plasmodium falciparum,* the parasite responsible for nearly all cases of severe malaria, must survive challenging environments to persist in its human host. Symptomatic malaria is characterized by periodic fevers corresponding to the 48-hour asexual reproduction of *P. falciparum* in red blood cells. As a result, *P. falciparum* has evolved a diverse collection of heat shock proteins to mitigate the stresses induced by temperature shifts. Among the assortment of heat shock proteins in *P. falciparum*, there is only one predicted canonical cytosolic J domain protein, HSP40 (PF3D7_1437900). Here, we generate a HSP40 tunable knockdown strain of *P. falciparum* to investigate the biological function of HSP40 during the intraerythrocytic lifecycle. We determine that HSP40 is required for malaria parasite asexual replication and survival of febrile temperatures. Previous reports have connected proteotoxic and thermal stress responses in malaria parasites. However, we find HSP40 has a specific role in heat shock survival and does not mitigate the proteotoxic stresses induced by artemisinin or proteosome inhibition. Following HSP40 knockdown, malaria parasites have a cell cycle progression defect and reduced nuclear replication. Untargeted proteomics reveal HSP40 depletion leads to a multifaceted downregulation of DNA replication and repair pathways. Additionally, we find HSP40 knockdown sensitizes parasites to DNA replication inhibition. Overall, these studies define the specialized role of the J domain protein HSP40 in malaria parasites during the blood stages of infection.

**Author Summary:** Malaria parasites have evolved to survive and cause disease in humans even when they are challenged by the temperature shifts of fever. We are interested in uncovering how malaria parasites persist despite exposure to febrile temperatures. We know the parasite has a diverse collection of heat shock proteins that are important for proper folding of many proteins within cells during stress conditions. In this study, we define the role of one specific heat shock protein, HSP40, by generating a strain of the human malaria parasite *P. falciparum* where we control the expression of HSP40 during the red blood cell stages of infection. We find that HSP40 is required for malaria parasites to replicate in red blood cells. We demonstrate HSP40 expression to be vital for malaria parasite survival of febrile temperature stress. Additionally, we determine that without HSP40, DNA replication and repair is disrupted. Our work has uncovered essential parasite biology that may be exploited for the development of new antimalarials.

## Introduction

Malaria is a persistent global health threat, causing over six hundred thousand deaths annually [1]. *Plasmodium* spp. are the intracellular parasites responsible for malaria and *Plasmodium falciparum* is the deadliest of all human malaria species. Humans respond to *P. falciparum* infection with a cyclical fever response corresponding to the synchronized replication of parasites in red blood cells [2–4]. As a result, *P. falciparum* has evolved unique mechanisms to tolerate host-induced febrile temperatures [5–9]. Resistance to front-line artemisinin-based combination therapies impedes efforts to the treat malaria worldwide [1]. Recent work has shown mechanisms the parasite employs to protect itself from febrile temperatures also appear to be utilized by the parasite to survive artemisinin treatment [9,10]. Defining fundamental malaria parasite biology as it pertains to stress survival is crucial to inform antimalarial design and combat rising drug resistance.

Processes related to the apicoplast in *P. falciparum* have been shown to be involved in the ability of malaria parasites to survive febrile temperatures [9,11]. The apicoplast houses the MEP pathway which generates isopentyl pyrophosphate (IPP), the building block for large isoprenoids that post-translationally modify proteins via prenylation [12]. Our previous work has shown that protein prenylation, specifically farnesylation, is immediately required for *P. falciparum* to survive temperature shifts like those experienced by the parasite during fever [11]. *P. falciparum* has a modest set of only four farnesylated proteins—one of these is the canonical J domain protein HSP40 (PF3D7_1437900), which has yet to be functionally characterized for its biological role in parasite growth and thermotolerance [13,14].

Molecular chaperones are crucial for folding nascent peptides and maintaining protein integrity during stress conditions [15]. J domain proteins load peptide substrates onto Hsp70 chaperones and mediate protein refolding by stimulating the ATPase activity of Hsp70 [16]. The diverse collection of J domain proteins guide substrate specificity for the restricted number of encoded Hsp70 chaperones [17–19]. *P. falciparum* has an expanded set of J domain proteins indicating the parasite has specifically evolved this class of chaperones for surviving the unique facets of the lifecycle [8]. In *P. falciparum,* HSP70-1 (PF3D7_0818900) partners with the J domain protein HSP40 and is required for parasite heat shock recovery; however, the role of HSP40 in thermotolerance remains undetermined [11,20–22].

In this study, we use a conditional knockdown approach to investigate the biological role of HSP40 in *P. falciparum* asexual replication, sensitivity to artemisinin, and thermotolerance. We find HSP40 is an essential protein for replication of malaria parasites in red blood cells and is vital for heat shock recovery. Additionally, we determine HSP40 is not involved in mitigating the proteotoxic stresses induced by artemisinin or proteosome inhibition. Interestingly, we find HSP40 depletion is associated with a multifaceted downregulation of DNA replication and repair pathways as well as an increased sensitivity to DNA replication inhibition. Altogether, this work teases apart the specialized role of HSP40 and uncovers unique biology as it pertains to thermotolerance and DNA replication in malaria parasites.

## Results

### HSP40 is an essential protein for *P. falciparum* asexual replication

Forward genetic screens and the inability to generate a HSP40 knockout strain suggest HSP40 is an essential gene in blood-stage *P. falciparum* [11,23]. Therefore, to investigate the biological function of HSP40 in asexual parasites, we employed the TetR-DOZI conditional knockdown system to control expression of HSP40 [24]. Due to regulation of the TetR-DOZI fusion protein and hairpin aptamers added to the mRNA of HSP40, HSP40 is expressed when anhydrotetracycline (aTc) is added to the culture media but substantially reduced when aTc is removed (Fig 1A). Using a CRISPR/Cas9 genomic editing strategy, the native locus of HSP40 was replaced with necessary components for TetR-DOZI regulation and confirmed by PCR amplification from genomic DNA (S1 Fig A). Upon removal of aTc, immunoblotting parasite lysates of the TetR-DOZI regulated HSP40 knockdown strain (hereafter referred to as HSP40^KD^) show reduced HSP40 expression beginning one day without aTc, confirming that TetR-DOZI regulates HSP40 as expected (Fig 1B).

**Fig 1.**
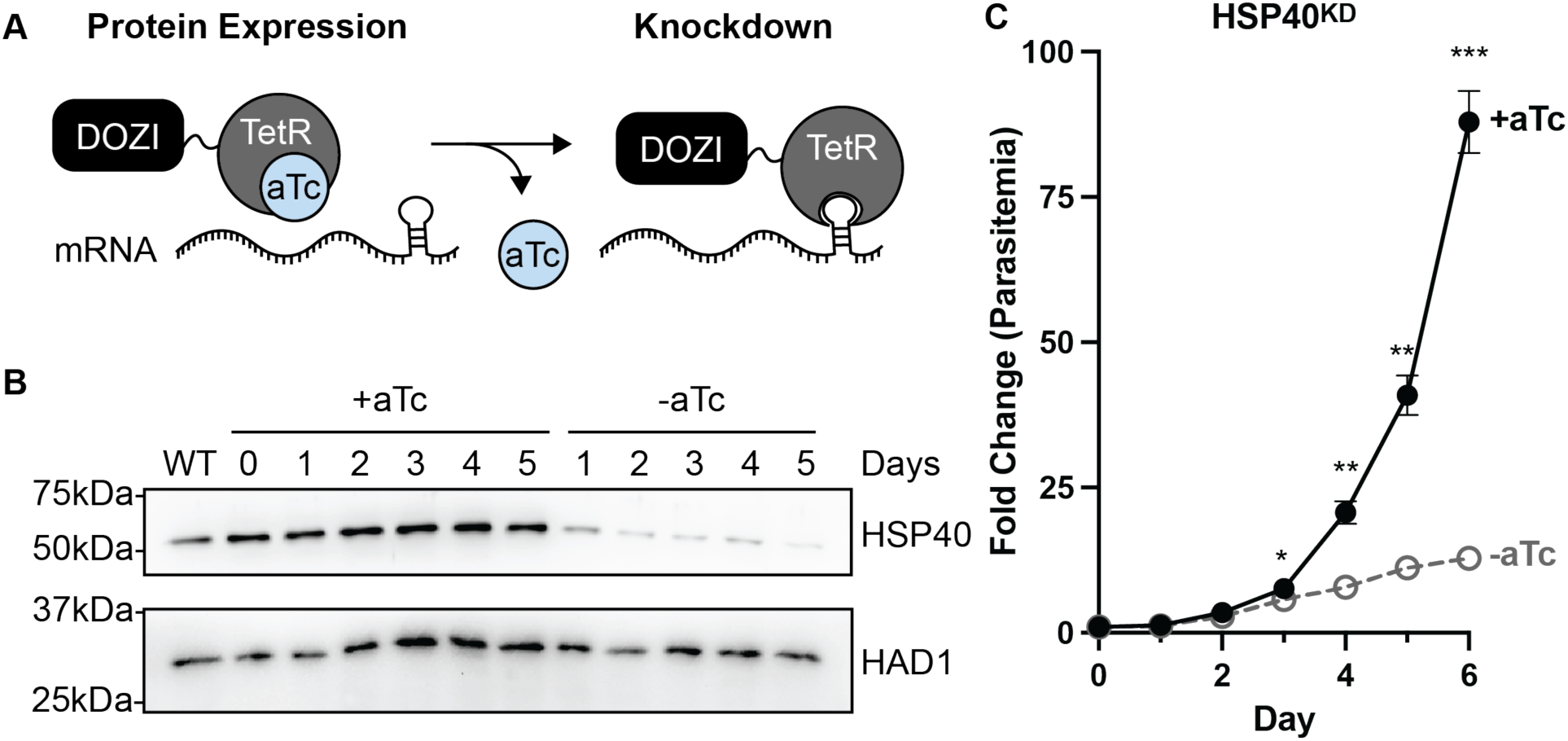
HSP40 expression is essential for *P. falciparum* replication. A) The TetR-DOZI system controls protein expression. Adding hairpin aptamers to the mRNA of the gene of interest allows for knockdown by removal of anhydrotetracycline (aTc) from culture media. B) Representative anti-HSP40 immunoblots of parasite lysates collected from HSP40^KD^ parasites +/− aTc for 5 days. HAD1 is used as a loading control. Without aTc, HSP40 expression decreases starting at 24 hrs. Blot is representative of three biological replicates. C) Growth assay of asynchronous HSP40^KD^ parasites. Fold change in parasitemia (percentage infected erythrocytes) quantified by flow cytometry every 24 hours from cultures grown +/− aTc. Parasites were split 1:6 after day 4. Data represents the mean +/− SEM of biological replicates, missing error bars are too small to be visualized. Parametric unpaired t-tests were performed (*p<0.05, **p<0.01, *** p<0.001).

To evaluate whether HSP40 plays a role in asexual replication of *P. falciparum*, we quantified asynchronous parasite growth over time after HSP40 knockdown. We find that HSP40 expression is essential for asexual replication of malaria parasites (Fig 1C). Interestingly, reduced HSP40 expression does not lead to a significant growth difference until after the first full 48-hour cycle of replication. We next examined whether the HSP40 knockdown effect was irreversible by adding aTc back to parasite cultures after two or four days without aTc (S1 Fig B-C). Adding aTc back to the cultures after two days without aTc, despite substantial reduction of HSP40 expression, parasites continue to grow with normal kinetics. Replenishing aTc after four days without aTc, when significant parasite growth delay has already occurred, parasites begin to replicate again. These results demonstrate that HSP40 knockdown results in substantial reduction of parasite replication but does not trigger irreversible cell death.

### HSP40 function is disrupted by N-terminal tagging

We next sought to generate pseudodiploid parasite strains that express an N-terminally tagged wildtype HSP40 in the HSP40^KD^ background. This would be an ideal tool for investigating the ability of variant forms of HSP40 to rescue knockdown phenotypes, to further define HSP40 domain-specific functions. Two independent transgenic parasite lines were generated with an N-terminal GFP- or FLAG tagged-HSP40 in the HSP40^KD^ background. While we confirm expression of the GFP- or FLAG-tagged constructs, neither construct rescue the growth defect of HSP40 knockdown (S2 Fig). Our data suggests that an N-terminal tag, even a small tag like FLAG, rendered HSP40 defective for its function. A C-terminal marker for the protein is not an option because HSP40 is post-translationally modified by prenylation which leads to cleavage of the C-terminal residues [25]. Using a complementation strategy to investigate the ability of HSP40 mutants to rescue the knockdown effect are beyond the scope of this inquiry.

### HSP40 knockdown sensitizes *P. falciparum* to heat shock

HSP40 is one of only four proteins that are modified post-translationally by farnesylation in *P. falciparum* [13,14]. Previous work has shown that protein farnesylation is required for malaria parasite thermotolerance [11]. We were able to take advantage of the delayed growth defect phenotype of the HSP40^KD^ parasites to determine if HSP40 plays a role in parasite heat shock recovery. Monitoring parasite growth after a 6-hour 41°C heat shock during the second day of HSP40 knockdown, we find that HSP40 depletion sensitizes parasites to heat stress (Fig 2). A similar but more modest effect is observed by heat shock on the first day of HSP40 knockdown, possibly mitigated by residual HSP40 (S3 Fig). These data demonstrate that HSP40 is essential for malaria parasites to survive fever-relevant heat shock temperatures.

**Fig 2.**
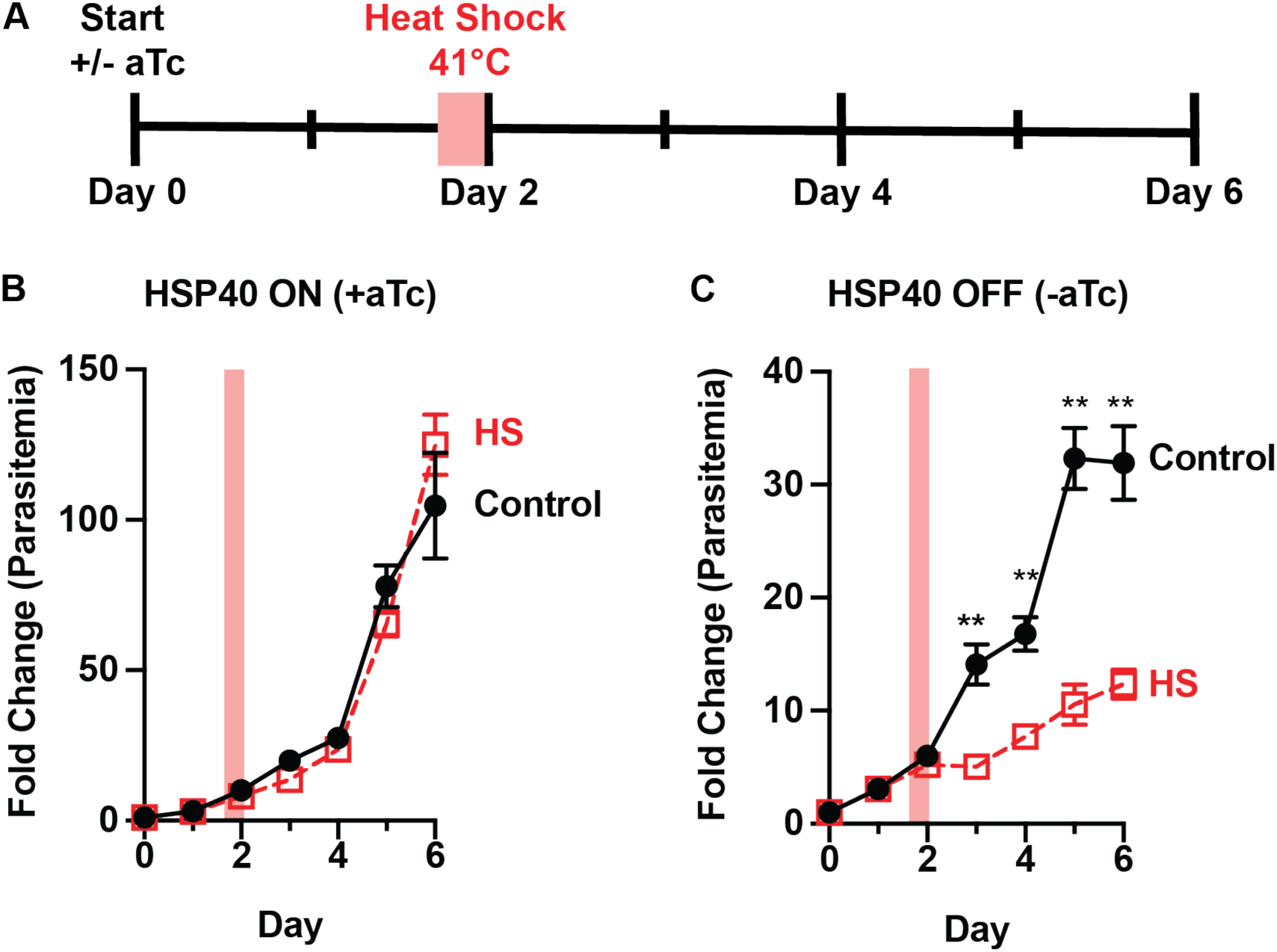
HSP40 expression is vital for *P. falciparum* survival following heat stress. A) Experimental design to assay thermotolerance: HSP40^KD^ parasites were subjected to a 6-hour 41°C heat shock (HS) on day two +/− aTc. Parasitemia was measured by flow cytometry collecting every 24 hours. Cultures were split 1:6 after day 4 collection. Growth assays measuring HSP40^KD^ after a 6hr heat shock show parasites B) recover from heat shock when HSP40 expression is on but C) lose this thermotolerance when HSP40 expression is off. Data represents the mean +/−SEM of three biological replicates, missing error bars are too small to visualize. Parametric unpaired t-tests were performed (**p<0.01).

### The HSP40 phenotype is partially rescued by growth at lower temperatures

While the temporary heat exposure showed HSP40 plays a role in heat shock recovery, we sought to evaluate whether HSP40 knockdown affects growth under constant exposure to different temperature conditions. Using a range of temperatures from 35°C to 38.5°C, we find that the requirement of HSP40 for parasite replication is temperature dependent (Fig 3). Growing parasites at 38.5°C, HSP40 knockdown leads to reduced parasitemia earlier than what is observed at 37°C (Fig 3B-C). Interestingly, at 35°C parasite replication does not show a significant reduction with HSP40 knockdown (Fig 3A). These results seemingly show a dose-effect, such that reduced HSP40 expression has a greater inhibitory effect on parasite replication with increasing temperature (Fig 3D-E). This demonstrates that not only is HSP40 expression critical for survival of fever-relevant heat pulses, but HSP40 also is an essential regulator of parasite growth under constant elevated temperature.

**Fig 3.**
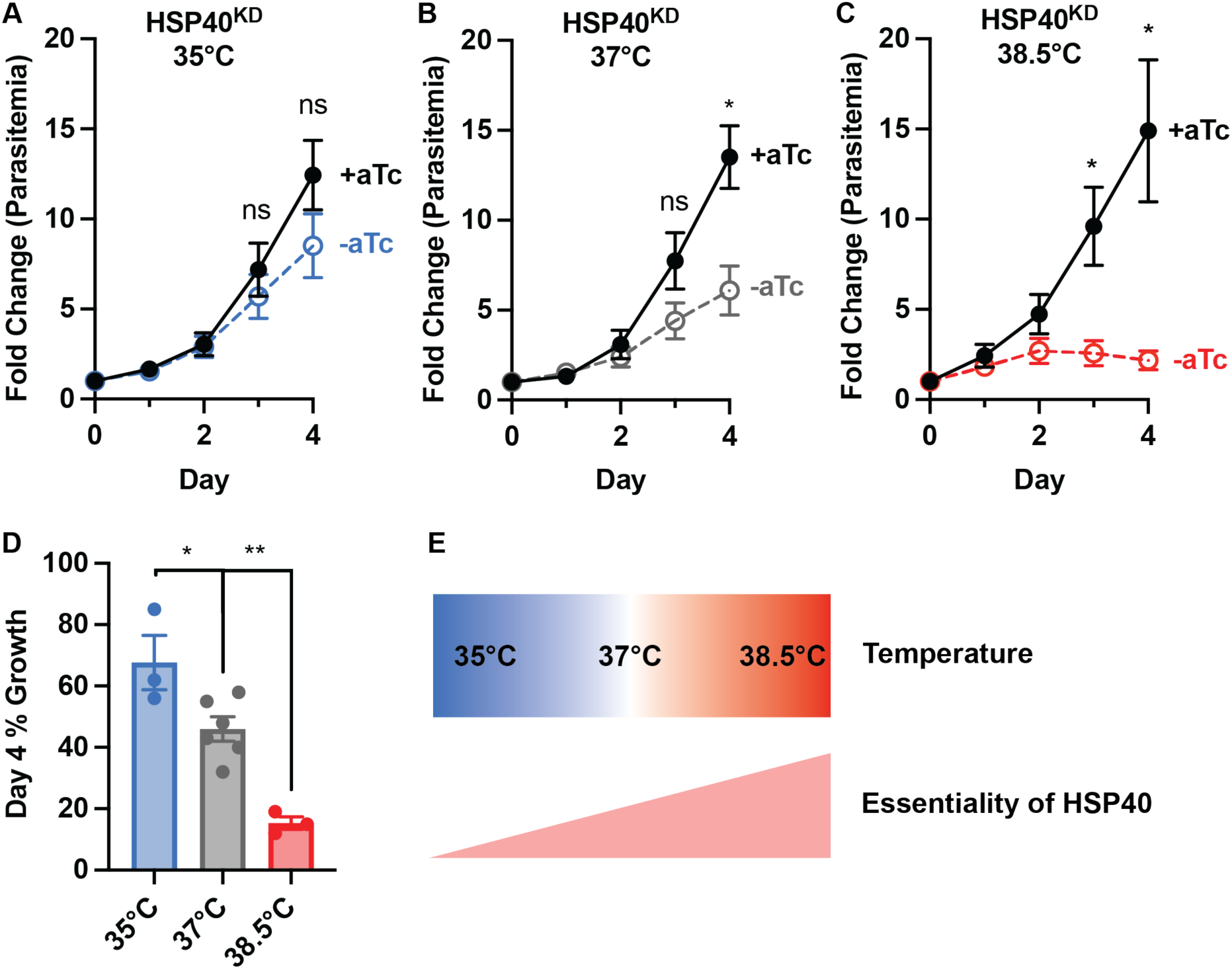
The HSP40 phenotype is partially rescued at lower temperatures. Growth assay of asynchronous HSP40^KD^ parasites measuring parasitemia by flow cytometry every 24 hours cultured +/− aTc at A) 35°C B) 37°C and C) 38.5°C. D) Normalizing the fold change in parasitemia on day 4 to the +aTc condition shows the HSP40 knockdown effect becomes more pronounced as temperature increases. Data represents the mean +/− SEM of biological replicates. Parametric unpaired t-tests were performed (*p<0.05, **p<0.01). E) The essential role of HSP40 increases with rising temperature.

### HSP40 does not protect parasites against artemisinin or proteosome inhibition

The antimalarial mechanism of artemisinin is, in part, due to an accumulation of damaged proteins [26]. Mechanisms of protection utilized by malaria parasites in heat shock recovery have been harnessed by the parasite to survive treatment with artemisinin [9,10]. Therefore, we measured sensitivity to the artemisinin derivative dihydroartemisinin (DHA) in HSP40^KD^ parasites +/−aTc to determine whether HSP40 mitigates proteotoxic stresses induced by this inhibitor. Even after two days of HSP40 depletion, we find no increased sensitivity to DHA. This phenotype was observed both under constant DHA treatment in asynchronous parasites (Fig 4A,D) as well as for a 6-hour DHA pulse against 0-3 hour ring stage parasites (RSA_0-3_; Fig 4B,E). To determine if phenotype was specific to DHA or applicable to other forms of chemically induced proteotoxic stress, we tested HSP40^KD^ sensitivity to bortezomib (BTZ), a potent proteosome inhibitor, with similar results (Fig 4C,F) [27]. Our findings highlight that the cellular response to temperature and proteotoxic stresses have distinct features. HSP40 has a specific role in heat stress survival and is not involved in mitigating the proteotoxic stresses induced by artemisinin or proteosome inhibition.

**Fig 4.**
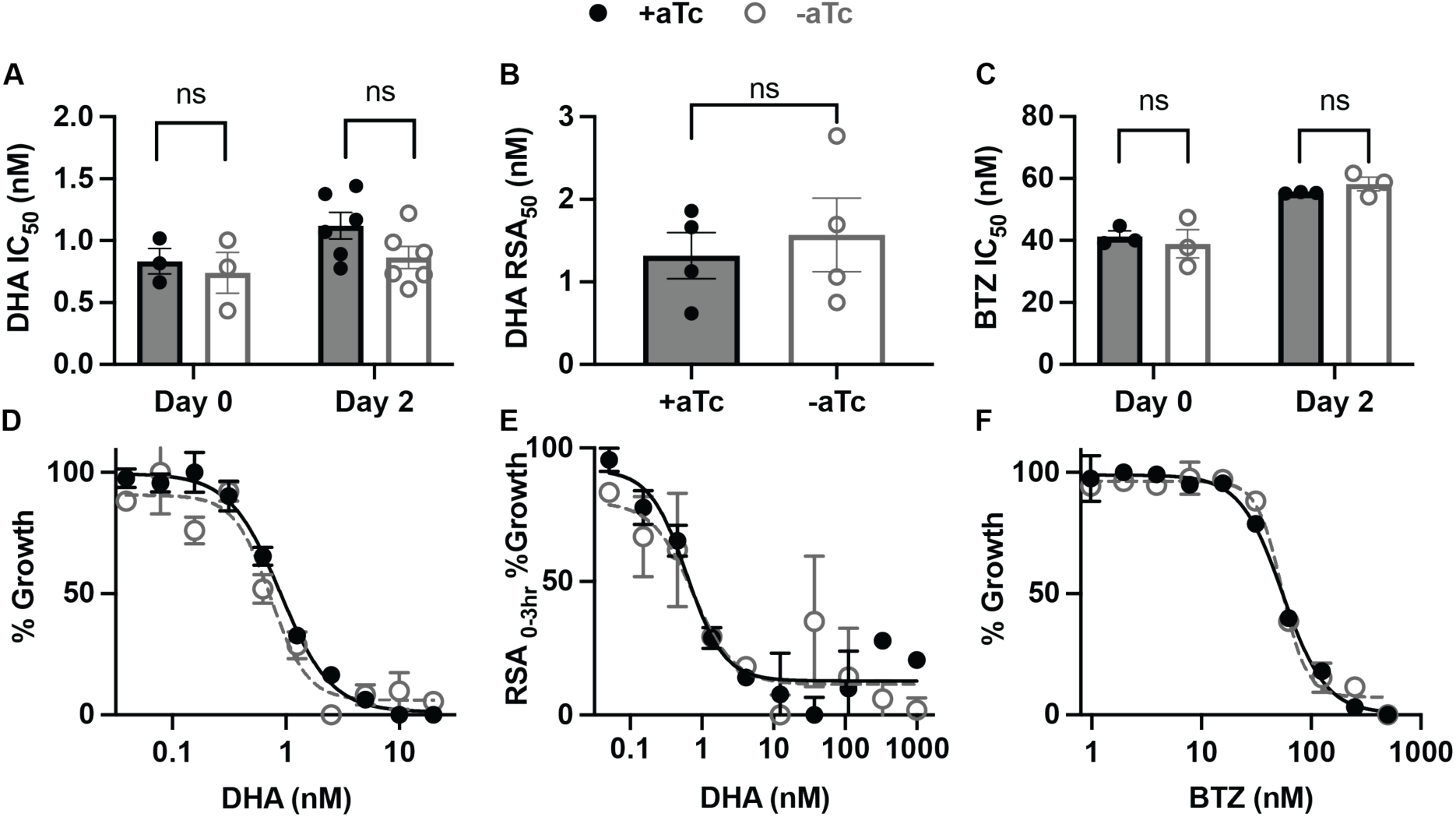
HSP40 does not mediate survival to chemically induced proteotoxic stress. A) Half-maximal inhibitory concentration (IC_50_) values of 72-hour dose-response curves started at day 0 or day 2 +/−aTc of asynchronous HSP40^KD^ parasites with dihydroartemisinin (DHA). B) DHA concentration value resulting in 50% growth inhibition during Ring-Stage Survival Assays (RSA_50_). A 6-hr DHA pulse at varying concentrations was given to 0-3 hour ring stage parasites on day 2 +/− aTc, parasite growth measured after 72 hours. C) IC_50_ summary values of 72-hour dose-response curves started at day 0 or day 2 +/−aTc of asynchronous HSP40^KD^ parasites with proteosome inhibitor bortezomib (BTZ). D) Representative IC_50_ curve of day 2 +/− aTc DHA treatment. E) Representative RSA_50_ curve. F) Representative IC_50_ curve of day 2 +/− aTc BTZ treatment. A-C summary values are of biological replicates performed in technical duplicate, data represents the mean +/−SEM. D-F are representative dose-response curves for one of the biological replicates showing the mean +/− SEM of the technical duplicates. No significance was found for the summary values performing parametric unpaired t-tests.

### HSP40 knockdown leads to a cell cycle progression defect and reduced nuclei replication

Having established HSP40 as an essential protein for parasite growth and thermotolerance, we next pursued deeper understanding of the mechanism by which decreased HSP40 expression leads to reduced parasite replication in red blood cells. We monitored lifecycle progression in tightly synchronized HSP40^KD^ parasites with and without aTc by microscopy as well as flow cytometry (Fig 5). Measuring fold change in parasitemia, we find that there is no difference in parasitemia until entering the third replication cycle (Fig 5A). Examining lifecycle progression by light microscopy reveals a developmental lag during the second cycle, beginning during parasite schizogony (S4 Fig A). In the second cycle, the HSP40 knockdown (-aTc) condition slows during late schizogony, eventually entering the third cycle, approximately 8 hours behind and with fewer parasites. The DNA content of these cells (measured by flow cytometry) highlights the lag in parasite development during the second cycle, as the peak in DNA content -aTc is approximately 8 hours behind the +aTc condition (Fig 5B). Additionally, throughout the entire second cycle the HSP40 knockdown parasites have reduced DNA content (Fig 5B).

**Fig 5.**
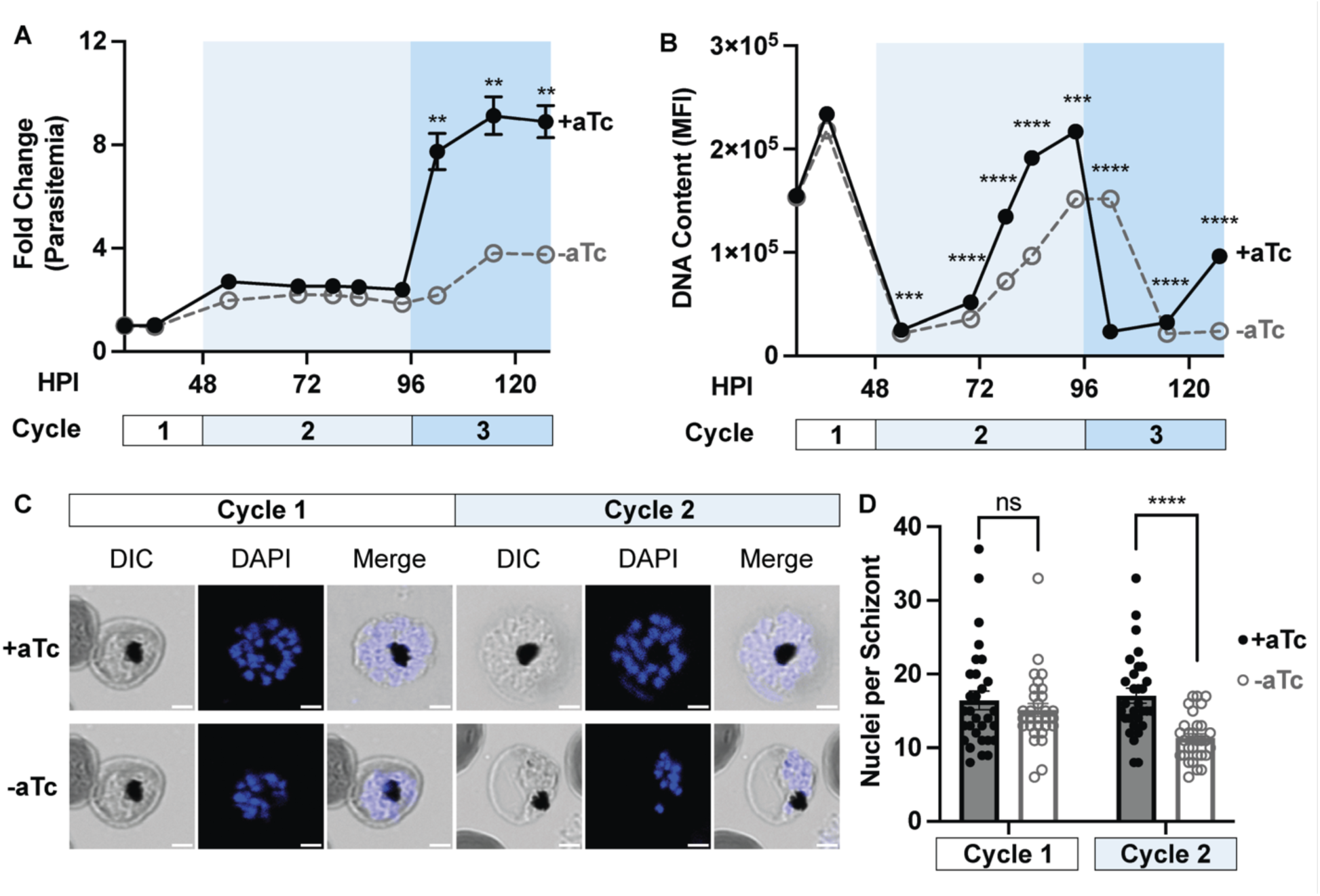
HSP40 depletion leads to a delayed reduction in parasite DNA content and nuclei generation. Tightly synchronized HSP40^KD^ parasites were starting +/−aTc at 8 hours post invasion in (HPI) and monitored through the third cycle of replication. A) Calculating fold change in parasitemia via flow cytometry shows loss in parasitemia entering the third cycle of HSP40 knockdown (-aTc). B) The median fluorescence intensity (MFI) of the DNA content of infected erythrocytes measured by flow cytometry demonstrates HSP40 knockdown corresponds to reduced level of DNA content throughout cycle 2. Additionally, the peak in DNA content is approximately 8 hours behind the +aTc condition. Data represents the mean +/− SEM of three biological replicates, missing error bars are too small to visualize. Parametric, unpaired t-test were performed (**p<0.01, ***p<0.001, ****p<0.0001). C) Representative image of the DAPI stained segmented schizonts used to quantify nuclei in the first and second cycle of HSP40 knockdown, scale bar represents 2 microns. D) Nuclei of segmented schizonts +/− aTc were counted for N=30 cells across three biological replicates in the first two replication cycles. Parametric, unpaired t-test were performed (****p<0.0001).

To evaluate whether these findings represented a defect in nuclear replication, we quantified the number of nuclei in segmented schizonts during the first two cycles of HSP40 knockdown. We find that HSP40 knockdown corresponded to a reduced number of nuclei per schizont in the second replication cycle (Fig 5C-D). Altogether these results show that HSP40 depletion leads to a delayed cell cycle progression defect during the trophozoite to schizont transition, reduced DNA content, and nuclear replication deficiency, ultimately resulting in fewer parasites.

### Downregulation of DNA replication and repair proteins following HSP40 depletion

Because HSP40 is predicted to have a role in maintaining protein homeostasis, we hypothesized that the nuclear replication defect we observed might be due to changes in protein expression during HSP40 knockdown [11,21,22]. For this reason, we performed whole-cell proteomics on trophozoites in cycle one and two of HSP40 knockdown. Consistent with the lack of phenotype in the first cycle +/− aTc, HSP40 is the only protein with significantly different expression at this time point (S5 Fig, S1 Table). This reduction in the abundance of HSP40 demonstrates that the TetR-DOZI regulation of HSP40 begins within the first cycle of aTc removal, but there are yet to be notable changes of any other protein within these parasites. In contrast, in the second cycle of HSP40 depletion we detect a total of 75 proteins with significantly different abundances, 67 downregulated and 8 upregulated proteins (Fig 6A, S2 Table). HSP40 is the most significantly downregulated protein among these parasites. Performing hierarchical clustering of the 75 hits reveals 14 other proteins regulated similarly to HSP40 across our 5 biological replicates (Fig 6C).

**Fig 6.**
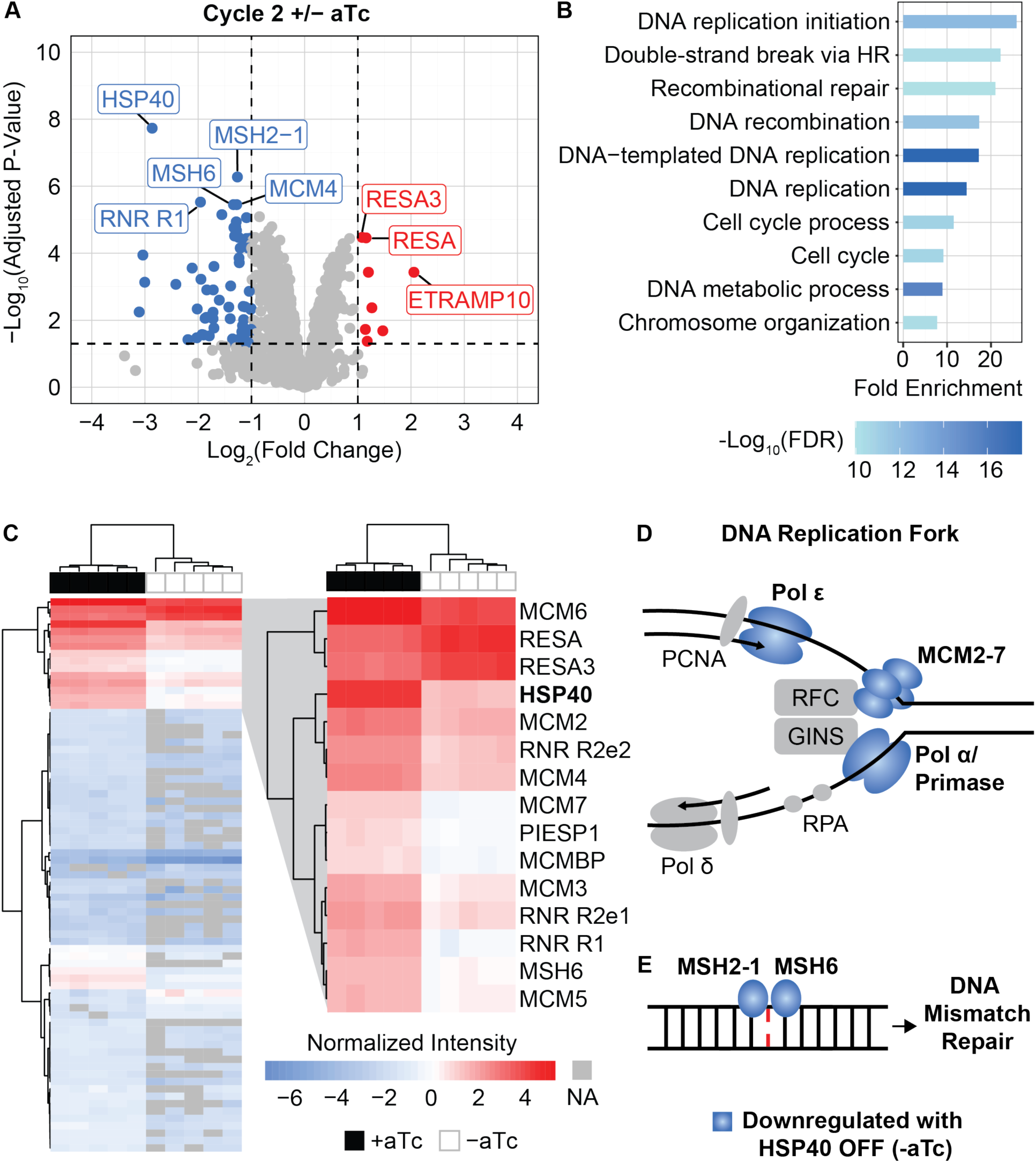
Proteomics reveal downregulation of DNA replication and repair pathways upon HSP40 knockdown. Synchronized HSP40^KD^ cultures were started +/− aTc as rings and collected for proteomics as trophozoites in cycles 1 and 2 of knockdown for N=5 biological replicates. A) Volcano plot of cycle 2 +/− aTc differential abundance analysis, downregulated genes in blue, upregulated in red. Differential abundance analysis was performed comparing +/− aTc for each cycle using LIMMA with empirical Bayes smoothing and Benjamini-Hochberg method for multiple test corrections. Significant hits had an adjusted P-value < 0.05 and absolute log_2_(fold change) > 1. B) Biological process gene ontology of the 67 downregulated proteins using Shiny GO 8.0. C) Heat map of the normalized intensity of all 75 differentially expressed proteins in cycle 2 +/− aTc across N=5 biological replicates detected by proteomics. Hierarchical clustering was performed using Euclidean distance and Ward method for columns and rows. Peptides that were not detected are NA in grey. The top three clusters which include HSP40 are zoomed in. Fully annotated heatmap shown in S5 Fig. D) Of the predicted proteins present at the DNA replication fork of *P. falciparum* during asexual replication, in cycle 2 of HSP40^KD^ -aTc we detect reduced abundances of the full MCM complex (MCM2-7), DNA polymerase alpha (Polα) subunit B, DNA primase large and small subunits, and DNA polymerase epsilon (Polε) subunit B. E) MSH2-1 and MSH6 detect DNA mismatches at the beginning of the mismatch repair pathway in *P. falciparum* and both of these have reduced protein abundance in cycle 2 of HSP40 knockdown.

Gene ontology analysis of the 67 downregulated genes reveals that proteins in DNA replication and repair pathways are affected by HSP40 knockdown (Fig 6B, S3 Table). We detect reduced expression of all six of the mini chromosome maintenance proteins (MCM 2-7). MCM2-7 drive formation of DNA pre-replication complexes as part of DNA replication licensing and comprise the helicase involved in DNA unwinding during replication (Fig 6D) [28,29]. In addition to the MCM complex, we detect downregulation of the DNA polymerase alpha (Polα) subunit B (PF3D7_1463300), DNA primase large (PF3D7_1438700) and small subunits (PF3D7_0910900), and DNA polymerase epsilon (Polε) subunit B (PF3D7_1234300), which comprise many of the known components involved in DNA replication forks in *P. falciparum* (Fig 6D) [30,31]. Additionally, we see reduced levels of MSH2-1 and MSH6, which work together to detect DNA mismatches in the initial steps of the mismatch repair pathway of *P. falciparum* (Fig 6E) [32]. These proteomics results demonstrate that HSP40 is required for homeostasis of DNA replication and repair machinery in malaria parasites.

### HSP40^KD^ parasites are hypersensitized to inhibition of DNA replication

Our proteomics indicated that DNA replication and repair pathways are markedly disrupted in during HSP40 knockdown. Clofarabine is a nucleoside analog drug that inhibits DNA replication machinery such as polymerases and primases, while simultaneously inducing DNA damage (Fig 7A) [33,34]. We find that HSP40 knockdown corresponds with increased sensitivity to clofarabine, indicating HSP40^KD^ parasites are hypersensitized to DNA replication inhibition (Fig 7B-C). This functionally validates our proteomics findings and highlights the dysregulation of DNA replication and repair during HSP40 knockdown.

**Fig 7.**
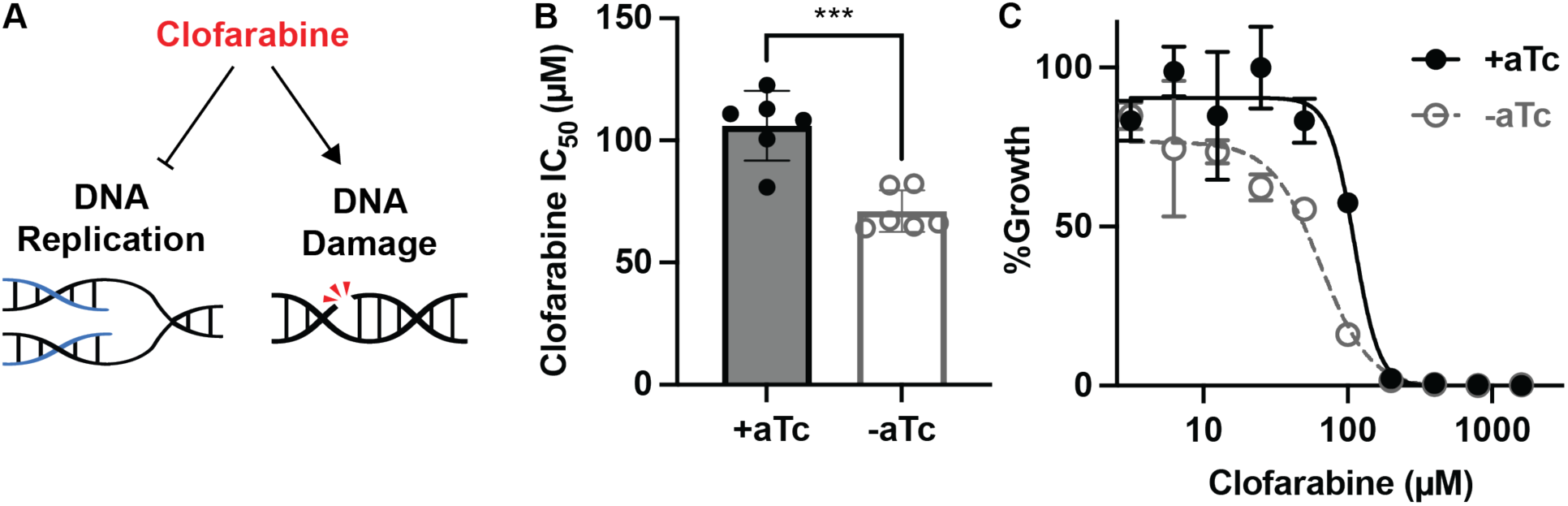
HSP40 knockdown sensitizes parasites to DNA replication inhibition. A) Clofarabine is a nucleoside analog which inhibits DNA replication while simultaneously eliciting DNA damage. B) IC_50_ summary values of 96-hour dose-response curves started with synchronous HSP40^KD^ ring parasites with clofarabine. Data represents the mean +/−SEM of N=6 biological replicates performed in technical duplicate. Parametric unpaired t-tests were performed (***p<0.001). C) Representative clofarabine IC_50_ curve for one of the biological replicates showing the mean +/− SEM of technical duplicates.

## Discussion

Replication of the malaria parasite within its human host requires the ability to tolerate modest temperature shifts, especially febrile temperatures > 38.0°C. Understanding malaria heat shock survival can define essential parasite biology and elucidate requirements for pathogenesis. The temperature changes during the malaria parasite lifecycle, combined with an unusually aggregation-prone proteome, have raised questions in the field as to how *P. falciparum* maintains protein homeostasis despite these challenges [35–37]. Molecular chaperones comprise a diverse class of proteins utilized in all biological systems to maintain protein integrity under standard and various stress conditions, including temperature stress. *P. falciparum* is thought to have a uniquely specialized collection of heat shock proteins that allow for protein stability in a temperature shifting environment [38,39]. In this work, we establish that the molecular chaperone, HSP40, is an essential regulator of *P. falciparum* blood stage replication and thermotolerance.

HSP40 belongs to the J domain protein family which comprises nearly 2 percent of the *P. falciparum* genome [40]. J domain proteins facilitate protein refolding by delivering misfolded peptides substrates to Hsp70 chaperones and stimulating Hsp70 ATPase activity [16]. The restricted number of Hsp70s in *P. falciparum* have their substrate specificity dictated by the wide assortment of J domain proteins [17–19]. HSP40 is classified as the canonical, cytosolic J domain protein and a fraction of HSP40 molecules are post-translationally modified by the isoprenyl group farnesyl [13,14,40]. Our previous work demonstrated that farnesylation is required for *P. falciparum* thermotolerance, and HSP40 is the sole farnesylated protein in *P. falciparum* with predicted roles in heat shock survival [11,13,14]. Interestingly, farnesylation of the HSP40 homolog in yeast and plants has been shown to be required for organismal thermotolerance, suggesting an evolutionarily conserved role [41–43]. Here we provided evidence that HSP40 has a specialized temperature-related essential function in malaria parasites. We find HSP40 is required for asexual parasite growth during both constant and temporary elevated temperature exposure; however, parasite growth is less dependent on HSP40 expression at lower temperatures. These results seemingly showed a dose-effect, such that reduced expression of HSP40 was more detrimental with increasing temperatures, highlighting the specificity in the essential function of HSP40 for heat stress survival.

Multiple studies have broadly connected intracellular processes that are essential for heat shock survival to parasite survival under treatment with the front-line antimalarial artemisinin. For example, heat stress and artemisinin induce similar transcriptional programs, including changes in expression of exported proteins and proteins involved in the unfolded protein response and lipid metabolism [44]. Heat-sensitive *P. falciparum* mutants tend to be sensitive to artemisinin and the proteosome inhibitor bortezomib [9]. Pre-treating malaria parasites with heat reduces their susceptibility to artemisinin, strongly connecting protective cellular processes under both stresses [10]. For these reasons, we were surprised to find that, although HSP40 is required for thermotolerance, it does not appear to play a role in mitigating the proteotoxic stresses caused by artemisinin derivative, dihydroartemisinin, or proteosome inhibitor, bortezomib. This could be due to distinct intracellular dysfunctions evoked by these stresses, requiring particular responses.

In *P. falciparum*, the specialized function of HSP40 appears to be related to homeostasis of DNA replication and repair machinery. During HSP40 knockdown parasites show a developmental lag during the DNA replication phase of the lifecycle, have a reduction in DNA content per cell and the number of nuclei per schizont. Our proteomics overwhelmingly point to DNA replication and repair pathways being downregulated upon HSP40 depletion. Finally, HSP40 knockdown corresponds with increased sensitivity to clofarabine which inhibits DNA replication and also elicits DNA damage. HSP40-mediated thermotolerance may be linked to its role in regulating the homeostasis of the DNA repair machinery. Malaria parasites upregulate DNA damage repair pathways and have detectable double stranded DNA breaks during heat shock [8,9,45]. The HSP40 homolog in yeast has been shown to be essential in regulation of DNA synthesis and damage responses, however the strategy by which HSP40 mitigates DNA replication and repair is unclear in *P. falciparum* [46]. DNA replication and repair downregulation during HSP40 knockdown suggests expression of these DNA protein components has a common mechanism of regulation. Understanding the mechanism by which reduced expression of HSP40 leads to disruption of DNA replication and repair machinery will likely yield additional insights into regulation of protein expression in *P. falciparum*.

In summary, this work defines the essential function of the J domain protein, HSP40, in *P. falciparum* asexual replication and thermotolerance. Our data tease apart the specialized role of HSP40, highlighting unique mechanisms malaria parasites have evolved to survive under different stress conditions. We identify HSP40 as a potential regulator of DNA replication and repair pathways, establishing a foundation for future inquiries in nuclear replication dynamics. Finally, this study elucidates vital parasite biology which could be exploited in the development on novel antimalarials and contributes to the broader understanding of this unique subclass of J domain molecular chaperones.

## Materials and Methods

### Parasite strains and cultures

Parasites were cultured in RPMI medium (Gibco) with the addition of 27mM NaHCO_3_, 11mM glucose, 5mM HEPES, 0.01mM thymidine, 1mM sodium pyruvate, 0.37mM hypoxanthine, 10ug/ml gentamicin, and 5g/L Albumax (Thermo Fisher Scientific) and maintained at 37°C in 5% O_2_, 5% CO_2_, 90% N_2_ in a 2% suspension of human red blood cells. The wild-type strain 3D7 (MRA-102) was obtained from BEI Resources Repository, NIAID, NIH. Deidentified red blood cells of either A+, AB+, or O+ blood type were obtained from the Children’s Hospital of Philadelphia Blood Bank and BioIVT. Parasites were synchronized using a combination of 5% sorbitol (Sigma: S889) and Percoll gradients (Sigma: P4927).

The HSP40 regulatable knockdown strain, HSP40^KD^, was generated by employing CRISPR/Cas9 to edit the native locus of HSP40 (Pf3D7_1437900) and incorporate TetR-DOZI regulation as previously described [47]. The pSN054 linear plasmid contained the segments that replaced the native locus by inserting aptamers at the 3’ end of the HSP40 gene as well as the TetR-DOZI fusion protein and the blasticidin selection marker. The pAIO3 vector contained the Cas9 enzyme along with the guide RNA to target the HSP40 locus. The pSN054 and pAIO3 vectors were generously provided to us from Daniel Goldberg’s lab. Plasmid sequences were confirmed via sanger sequencing.

For transfection, 50ug of each plasmid was precipitated and resuspended in 400uL of Cytomix (120mM KCl, 0.15mM CaCl_2_, 2mM EGTA, 5mM MgCl_2_, 10mM K_2_HPO_4_, 25mM HEPES, pH 7.6). Synchronized 3D7 ring parasites at roughly 5% parasitemia were washed with Cytomix and resuspended in the 400uL Cytomix-DNA mixture. Cells were electroporated at 950uF capacitance and 0.31kV using a Biorad Genepulser Xcell. Cells were washed with media and cultured immediately with the addition of 50-100nM aTc (Caymen Chemicals: 10009542) diluted in DMSO. Starting 24 hours post-transfection, parasites were selected for using 2.5 ug/mL blasticidin (Invitrogen: R210-01).

From the pooled population of transfected parasites, individual clonal populations were grown out. Five individual clones were isolated and their growth +/− aTc was characterized to have a similar phenotype. One of the clones was used for experiments in biological replicates, defined as cultures grown and treated separately for at least one week.

Genomic integration of the TetR-DOZI cassette was confirmed using test PCR as well as sequencing PCR fragments from genomic DNA from HSP40^KD^ clones. Genomic DNA was isolated using chloroform extraction and PCRs were performed using Primestar GXL DNA Polymerase (Takara: R050A) according to manufacturer’s instructions.

### Generating the complement HSP40 strains

To generate the pseudo-diploid HSP40 strains, the piggyBac transposon system was used to integrate a second copy of the HSP40 gene with either an N-terminal GFP or 3x FLAG tag into the HSP40^KD^ genome under WR92210 selection [48]. The pTEOE plasmid with a Hsp86 promoter housed the tagged HSP40 coding sequence. The pTEOE plasmid along with the pHTH helper plasmid were transfected together, then kept under 5nM WR92210 after 24 hours along with the blasticidin and aTc. As further confirmation, the FLAG-HSP40 insert was PCR amplified from genomic DNA and sequenced to confirm genomic integration.

### Western blotting

Parasite lysates were obtained by 1% saponin lysing 25mls of parasite cultures followed by cold PBS washes. Samples were stored in −80°C until ready to sonicate. All steps for sample preparation were performed at 4°C. Samples were washed in lysis buffer (10% glycerol, 100mM NaCl, 100mM Tris pH 7.5, 1mM MgCl_2_, 1mM DTT, with an EDTA-free protease inhibitor cocktail mini-tablet (Roche: 11836170001)) twice. Parasite pellets were resuspended in lysis buffer then sonicated with 6 cycles, 10 second pulses at 40% amplitude with a FisherBrand Model 120 Sonic Dismembrator. After sonication, samples were stored at −80°C. For western blotting, sonicated supernatant was diluted in SDS-PAGE buffer with 2-β-mercaptoethanol and boiled for 10min. The equivalent of approximately 1×10^7^ parasites were loaded on 12% SDS-PAGE gels and ran at 120V, transferred to a PVDF membrane using the BioRad Transblot Turbo system with TBT-0.05% SDS buffer, and blocked overnight with 5% BSA in PBS_T_ rocking at 4°C. Primary antibody and secondary antibody was incubated for an hour at room temperature rocking with three 10-minute PBS_T_ washes in between.

*P. falciparum* HSP40 rabbit anti-sera generated previously [11] was used at a 1:5000 dilution, *P. falciparum* HAD1 rabbit anti-sera generated previously [49] was used at a 1:20,000 dilution in 5% BSA in PBS_T_. Mouse anti-FLAG antibody (Sigma F1804) was used at a 1:1,000 in 3% Milk in PBS_T_. Secondary antibody goat anti-rabbit HRP (Thermofisher: 65-6120) and goat anti-mouse HRP (Thermofisher: 31430) were used at 1:20,000 in 5% BSA in PBST. Western blots were developed using SuperSignal West Pico Plus (Thermofisher: 34580) and imaged using a BioRad ChemiDoc.

### Growth assays

The growth assays +/− aTc were started by washing parasite cultures three times to ensure removal of aTc, and then diluted to 1% parasitemia. A negative empty red blood cell 2% hematocrit control was prepared and treated in parallel. Flow cytometry samples were collected every 24 hours. Cultures were given fresh media every two days or split as indicated to maintain healthy cultures.

### Heat shock assay

A growth assay was set up and analyzed as described above, however at the indicated timepoint cultures were moved to an incubator set to 41°C for 6 hours. Media was exchanged, and parasites were returned to 37°C for the remaining experiment.

### Cell cycle progression

To monitor lifecycle progression in tightly synchronized HSP40^KD^ parasites, cells were synchronized using a combination of sorbitol and percoll gradients. When parasites were approximately 8-hour rings, +/− aTc was started by washing parasite cultures three times to ensure removal of aTc, and then diluted to 2-3% parasitemia. Microscopy slides and flow cytometry samples were collected for multiple timepoints from the initial replication cycle all the way to the middle of cycle 3 to fully capture the cell cycle progression. Media was exchanged 10 times throughout the experiment to maintain healthy cultures.

### Flow cytometry

For flow cytometry samples, 50uL of cells were fixed and stored in 4% paraformaldehyde, 0.025% glutaraldehyde at 4°C. When ready to analyze, cells were washed with PBS and resuspended to 1% hematocrit in PBS. Then 50uL of cell suspension was diluted into 300uL of 0.3ug/mL acridine orange (Invitrogen: A3568) in PBS and analyzed by a Cytek Aurora flow cytometer. Gating and parasitemia was determined using FlowJo software gating for red blood cells (SSC-A vs FSC-A), single cells (SSC-A vs SSC-H), and then for infected red blood cells (B3-A vs B7-A). Fold-change in parasitemia was determined by subtracting the uninfected red blood cell control collected at the same time, dividing each sample first timepoint parasitemia, and then multiplying by the split factor if cultures were split. To measure DNA content, median fluorescence intensity of the infected red blood cell population for the B3-A was used.

### Light microscopy

Microscopy slides of synchronized parasites blood smears were fixed 10 seconds in methanol throughout the asexual parasite lifecycle +/− aTc. All fixed slides were stained for 15min with Giemsa stain (Sigma: SLCM6930) diluted 1:20 and imaged with an Olympus CX43 microscope and Olympus DL21 camera at 100X magnification. Images were cropped, adjusted for brightness, and exported using ImageJ.

### Fluorescence microscopy

#### Nuclei counting

Thin blood smears of late-stage HSP40^KD^ schizonts +/− aTc were prepared on Superfrost glass slides (Fisherbrand: 12-550-15) and air-dried. Smears were fixed in chilled methanol and stored at −20°C. During cycle 2, since HSP40^KD^ -aTc parasites lag in their progression, smears were made approximately 8 hours later than +aTc parasites. On the day of staining, the smears were air-dried and mounted with Prolong Gold Antifade with DAPI (Molecular probes: P36941). Images were captured using a Leica confocal DMi-8 microscope with a 40x/1.35 numerical aperture (NA) oil immersion objective. Serial z-sections of each image were gathered, and the z-stack with the best representation is illustrated in the figure. Images were analyzed by open-sourced ImageJ software. Approximately 30 individual schizonts in each parasite sample were scored using ImageJ thresholding combined with watershed and analyze particles.

#### GFP-HSP40

100uL of live parasites at approximately 5% parasitemia in 2% hematocrit were washed twice with HBSS, then stained with Hoechst 33258 (Invitrogen: H21491) at a 40ug/mL concentration for 20 minutes in the dark at 37°C. After staining, cells were washed twice with HBSS, then concentrated to 10% hematocrit and added to Superfrost slides (Fisherbrand: 12-550-15) with coverslips and imaged with a Nikon Eclipse Ti2 inverted microscope. Images were cropped and exported using ImageJ.

### Drug sensitivity assays

All dose-response inhibition experiments were performed on at least three biological replicates of HSP40^KD^ parasites in technical duplicate +/− aTc. Parasite growth was measured on a CLARIOstarPlus (BMG LAB TECH) plate reader with Quant-iT PicoGreen dsDNA reagent (Invitrogen: P7581) staining. Data were fit to a non-linear regression to determine 50% inhibitory concentration (IC_50_) value using GraphPad Prism.

Dihydroartemisinin (DHA) and Bortezomib (BTZ) IC_50_ experiments were done with asynchronous HSP40^KD^ parasites on the same day as removing aTc or after two days without aTc. DHA (Caymen Chemicals: 19846) and BTZ (Caymen Chemicals: 10008822) were diluted in DMSO. For DHA two-fold dilutions from 0nM to 20nM and for BTZ, two-fold dilutions from 0nM to 400nM were added to 100uL of parasites and analyzed after 72 hours.

DHA Ring-stage Survival Assays (RSA) were done with tightly synchronized 0-3hr HSP40^KD^ rings after two days +/− aTc, adapted from methods as described [50]. Three-fold dilutions from 0nM to 1uM of DHA were added to 100uL of parasites for 6hrs. After the 6-hour DHA pulse, media was exchanged four times and cells were transferred to a new plate, maintained with or without aTc and analyzed after 72 hours.

Clofarabine IC_50_ experiments were done with synchronous 8-hour ring HSP40^KD^ parasites on the same day as removing aTc. Clofarabine (Sigma: C7495) was diluted in DMSO and added in two-fold dilutions from 0uM to 1.6uM to 100uL of parasites analyzed after 96 hours.

### Proteomics

#### Sample preparation

Proteomics samples were obtained by tightly synchronizing HSP40^KD^ parasites and washing off aTc as rings. During the cycle 1 and cycle 2 a sample was collected +/− aTc for five biological replicates when parasites were predominately trophozoite stage. For each replicate at the two separate time points, a 30mL 4% hematocrit sample with at least 5% parasitemia was collected. Samples were prepared by washing with PBS, adding 1% saponin to lyse red blood cells, then washing twice with cold PBS prior to flash freezing on dry ice. Parasite pellets were stored at −80°C until ready to run on LC-MS/MS by the CHOP Proteomics Core.

#### In-solution digestion

Parasite pellets underwent lysis, solubilization, and digestion on an S-Trap (Protifi) following the manufacturer’s protocol [51]. Subsequently, the resulting peptides were de-salted using an Oasis HLB plate (Waters), dried via vacuum centrifugation, and reconstituted in 0.1% TFA containing iRT peptides (Biognosys Schlieren, Switzerland).

#### Mass spectrometry acquisition and data analysis

Peptides were analyzed on a QExactive HF mass spectrometer coupled with an Ultimate 3000 nano UPLC system and an EasySpray source utilizing data independent acquisition (DIA). Raw data were searched using Spectronaut [52]. The MS2 intensity values for proteins generated by Spectronaut were used for bioinformatics analysis. Proteomics data processing and statistical analysis were conducted in R. The MS2 intensity values generated by Spectronaut were utilized for analyzing the entire proteome dataset. The data underwent log2 transformation and normalization by subtracting the median value for each sample. To ensure data integrity, we filtered it to retain only proteins with complete values in at least one treatment group (+ or - aTc). To compare proteomics data across groups, we employed a Limma (linear models for microarray data) t-test to identify proteins with differential abundance, with empirical Bayes smoothing and Benjamini-Hochberg method for multiple test correction. Lists of differentially abundant proteins were generated based on criteria of adjusted P. value < 0.05 and absolute log2 fold change greater than one, resulting in a prioritized list for subsequent analysis.

#### Gene ontology and heat map

Gene ontology (GO) analysis was done using ShinyGO, using the genes detected by proteomics as the background a false discovery rate cut-off of 0.05 and the GO Biological Process database [53]. For the heat map, Ward’s hierarchical clustering was done using Euclidean clustering distances for rows and columns. Data were processed and visualized using R studio.

## Acknowledgements

We thank Daniel Goldberg (Washington University of St. Louis) for supplying the pSN054 and pAIO3 plasmid vectors and the CHOP Proteomics Core facility for their assistance with our proteomics screen. This work was supported by the NIH/NIAID R01AI103280 (AOJ), R01AI171514 (AOJ), F32AI138373 (ESM), T32AI007172 (ESM), T32AI007532 (BR), the Pathogenesis of Infectious Diseases Burroughs Wellcome Fund, the Doris Duke Foundation Paragon of Research Excellence Award, and the Children’s Hospital of Philadelphia.

## Supporting Information

**S1 Fig.**
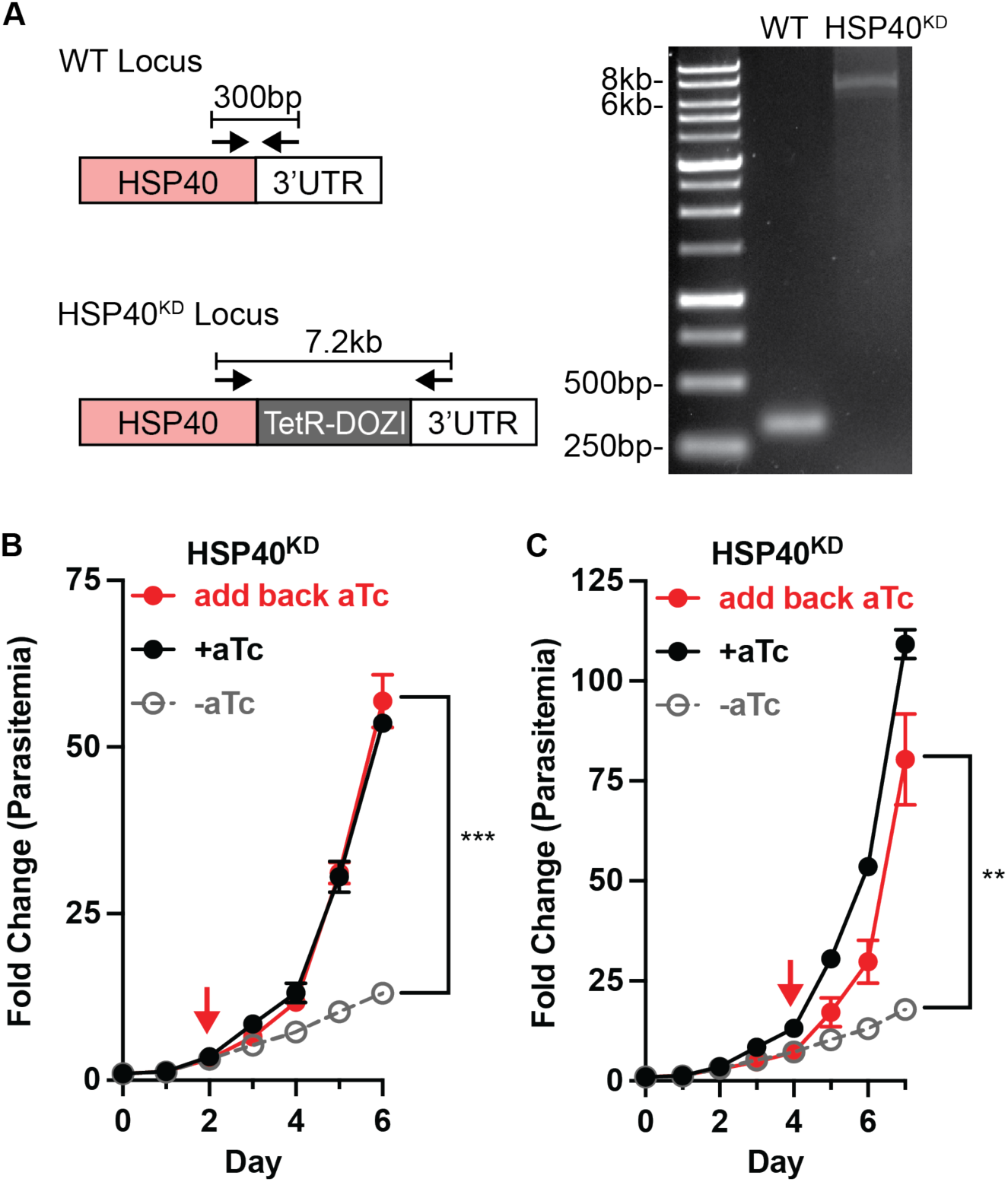
HSP40^KD^ parasites demonstrate a reversible knockdown phenotype. A) PCR tests confirm genomic integration of the TetR-DOZI cassette at the HSP40 locus in *P. falciparum.* The same primer set (indicated by black arrows) was used for PCR with 3D7 (WT) and HSP40^KD^ genomic DNA. Growth assays of asynchronous HSP40^KD^ parasites measuring fold change in parasitemia by flow cytometry every 24 hours cultured +/− aTc, adding back aTc on either B) Day 2 or C) Day 4 -aTc (indicated by red arrow). Parasites were split 1:6 after day 4. Data represents the mean +/− SEM of three biological replicates, missing error bars are too small to be visualized. Parametric unpaired t-tests between the add back aTc and -aTc condition were performed for the final day of collection (**p<0.01, *** p<0.001).

**S2 Fig.**
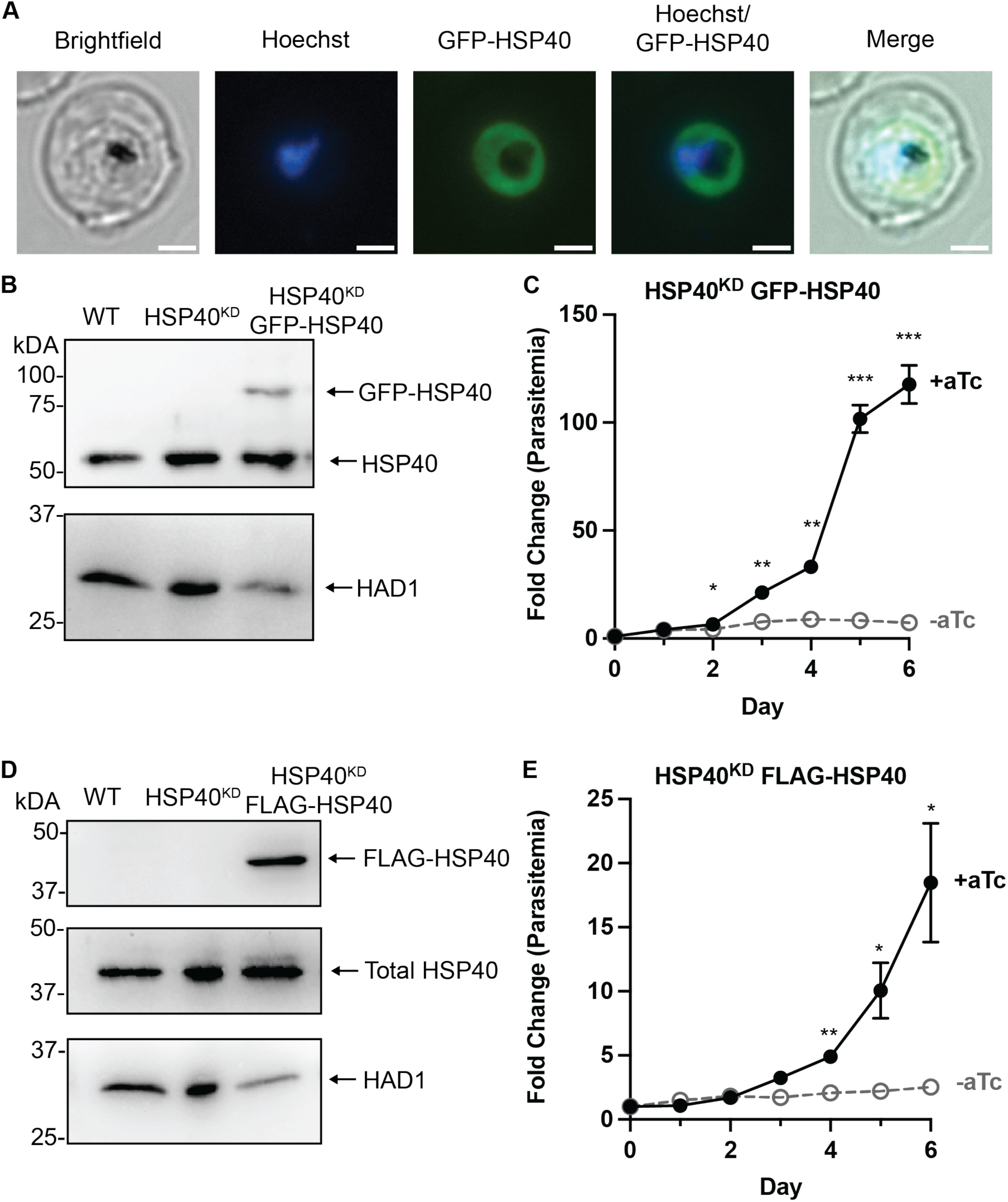
The function of HSP40 is disrupted by N-terminal tagging. A) Representative fluorescence microscopy image shows HSP40^KD^ GFP-HSP40 parasites express GFP-HSP40, scale bar represents 2 microns. B) Representative anti-HSP40 western blots of parasite lysates collected from HSP40^KD^ GFP-HSP40 parasites show expression of the GFP-HSP40 fusion protein. HAD1 was used as a loading control. Blot is representative of three biological replicates. C) Growth assay of asynchronous HSP40^KD^ GFP-HSP40 parasites measuring parasitemia by flow cytometry every 24 hours cultured +/− aTc. Parasites were split 1:6 after day 2. Data represents the mean +/− SEM of three biological replicates, missing error bars are too small to be visualized. Parametric unpaired t-tests were performed (*p<0.05, **p<0.01, *** p<0.001). D) Representative anti-FLAG and anti-HPS40 western blots of parasite lysates collected from HSP40^KD^ FLAG-HSP40 parasites show the complement stain expresses FLAG-HSP40. HAD1 was used as a loading control. Blot is representative of three biological replicates. E) Growth assay of asynchronous HSP40^KD^ FLAG-HSP40 parasites measuring parasitemia by flow cytometry every 24 hours cultured +/− aTc. Parasites were split 1:6 after day 4 collection. Data represents the mean +/− SEM of three biological replicates, missing error bars are too small to be visualized. Parametric unpaired t-tests were performed (*p<0.05, **p<0.01).

**S3 Fig.**
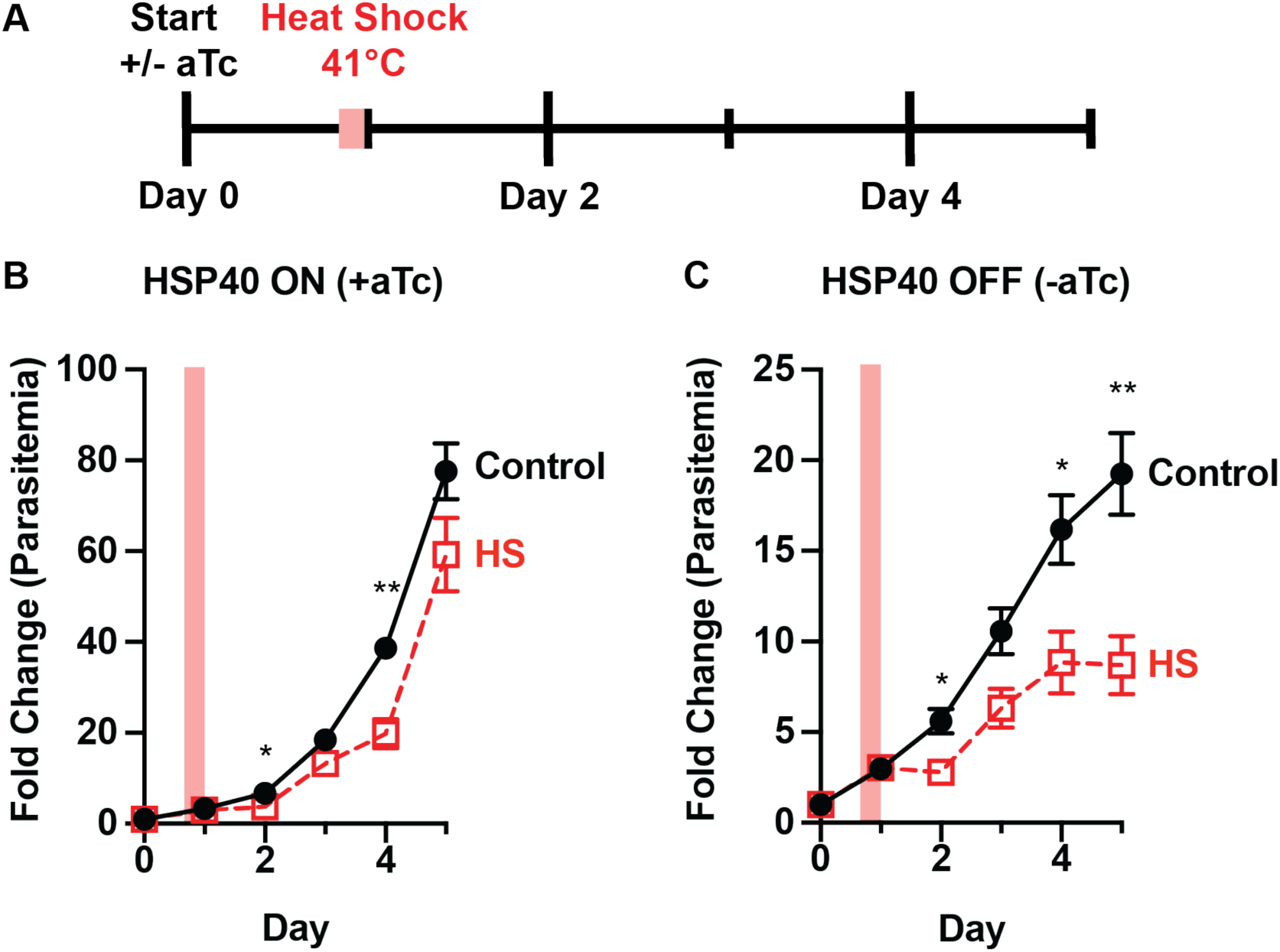
Heat shock after one day of HSP40 knockdown shows reduced parasite thermotolerance. A) Experimental design to assay thermotolerance: HSP40^KD^ parasites were subjected to a 6-hour 41°C heat shock (HS) on day one +/− aTc. Parasitemia was measured by flow cytometry collecting every 24 hours. Cultures were split 1:4 after day 3 collection. Growth assays measuring HSP40^KD^ after a 6hr heat shock (HS) show parasites recover from heat shock when B) HSP40 expression is on but lose this ability when C) HSP40 expression is off. Data represents the mean +/−SEM of biological replicates, missing error bars are too small to visualize. Parametric unpaired t-tests were performed (*p<0.05, **p<0.01).

**S4 Fig.**
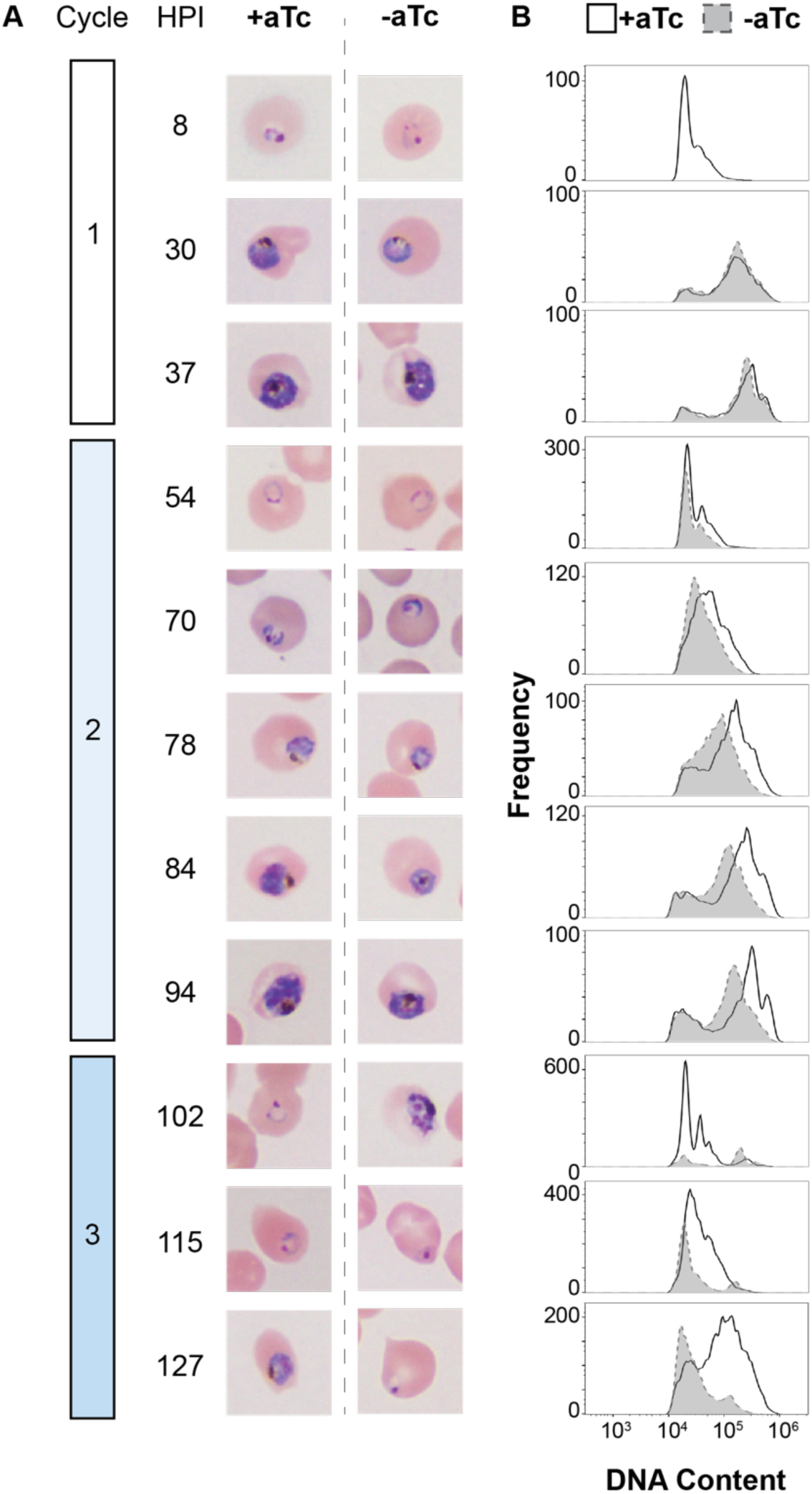
HSP40^KD^ parasites demonstrate a second cycle developmental defect. A) Tightly synchronized HSP40^KD^ parasites were monitored for lifecycle progression starting +/−aTc at 8 hours post invasion (HPI) through the third cycle of replication. During Cycle 2, there is a developmental lag starting when +aTc is 84hpi and continues as +aTc parasites enter cycle 3. Data is representative of 3 biological replicates. B) Histograms of infected red blood cell DNA content in HSP40^KD^ parasites +/− aTc from flow cytometry samples collected at time points indicated in part A. Starting at 84 HPI when the +aTc condition progresses into schizogony and increases the DNA content of cells, the -aTc condition lags. Entering cycle 3 at 102 HPI, the +aTc condition shows a large population with predominantly lower DNA content due to the newly invaded cycle 3 rings, while the -aTc has a smaller total population of cells with higher DNA content. Data is representative of 3 biological replicates.

**S5 Fig.**
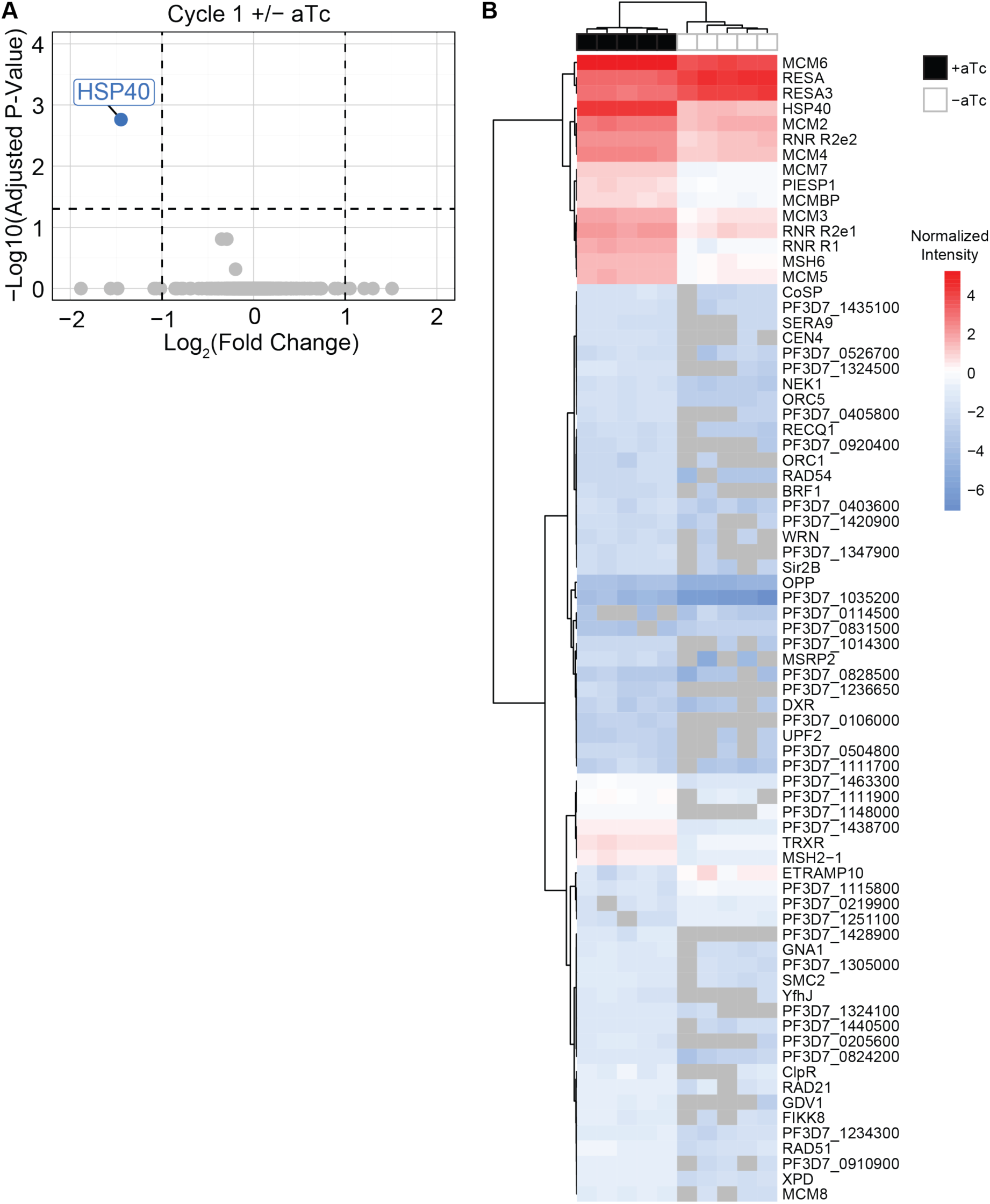
Whole-Cell Proteomics of HSP40^KD^ parasites +/−aTc. A) Volcano plot of cycle 1 +/− aTc differential abundance analysis, HSP40 was the only protein with significantly different expression. B) Heat map of the normalized intensity of all 75 differentially expressed proteins cycle 2 +/− aTc across N=5 biological replicates detected by proteomics. Hierarchical clustering was performed using Euclidean distance and Ward method for columns and rows. Peptides that were not detected are NA in grey.

**S1 Table. HSP40 Cycle 1 Proteomics**

**S2 Table. HSP40 Cycle 2 Proteomics**

**S3 Table. HSP40 Cycle 2 Downregulated GO Analysis**

